# Tissue-aware elastic net decomposition reveals shared and lineage-specific drug response biomarkers

**DOI:** 10.64898/2026.06.21.733619

**Authors:** Jared Strauch, Leila Azinfar, Heather H. Pua, James P. Long, Kevin R. Coombes, Amir Asiaee

**Affiliations:** Department of Biostatistics, Vanderbilt University Medical Center, Nashville, Tennessee, 37232, USA; Department of Pathology, Microbiology and Immunology, Vanderbilt University Medical Center, Nashville, Tennessee, 37232, USA; Department of Biostatistics, The University of Texas MD Anderson Cancer Center, Houston, Texas, 77030, USA; Department of Biostatistics, Data Science, and Epidemiology, Georgia Cancer Center, Augusta University, Augusta, Georgia, 30912, USA

**Keywords:** drug response prediction, elastic net, multitask learning, tissue specificity, data leakage, pharmacogenomics, interpretability

## Abstract

**Motivation:** Computational models that predict cancer drug response from genomic features are central to biomarker discovery, yet a recent audit found data leakage in 72% of 32 published methods, and complex models offer little interpretability while only modestly exceeding simple baselines under honest evaluation. Tissue lineage is a largely untapped source of legitimate inductive bias, but existing tissue-aware methods neither separate pan-cancer from lineage-specific signal nor report leakage-free performance.

**Results:** We introduce the Data Shared Elastic Net (DSEN), a tissue-aware regression that decomposes each drug’s model into a shared coefficient block common to all lineages and tissue-specific deviation blocks. Under leakage-free cross-validation across 265 drugs, 1,462 cell lines and 31 tissue lineages, DSEN improved mean squared error over a standard elastic net for 92.5% of drugs (mean 4.95%) while selecting 58% fewer stable shared features. Shared coefficients generalized to held-out tissues (59% tissue-level win rate) and recurrently recovered transferable pathway modules (p53, MAPK), whereas tissue blocks captured lineage markers such as the skin *MITF* /*S100B* program. The closest tissue-aware comparator, TG-LASSO, performed worse than the tissue-agnostic baseline (−13.8% mean MSE). Ablation shows tissue-aware modeling helps most when features are scarce, with no single modality dominating.

**Availability and implementation:** https://github.com/AsiaeeLab/tissue-aware-drug-response.

## Introduction

Genomic markers that predict drug sensitivity in cancer cells are routinely nominated using machine learning models trained on large-scale pharmacogenomic screens such as the Cancer Cell Line Encyclopedia (CCLE) and Genomics of Drug Sensitivity in Cancer (GDSC) [Barretina et al., 2012, Garnett et al., 2012, Iorio et al., 2016, Ghandi et al., 2019]. These models identify molecular features—gene expression, mutations, copy number alterations, and protein levels—that stratify cell lines by drug response and may ultimately guide patient selection in clinical trials [Costello et al., 2014, Geeleher et al., 2014]. Over the past decade, the field has grown rapidly, with more than 40 published methods spanning classical machine learning and deep learning architectures [Firoozbakht et al., 2022, Adam et al., 2020, Baptista et al., 2021].

Despite this proliferation, several recent studies have exposed fundamental concerns about the reliability and utility of reported results. A recent audit of 32 drug response prediction methods found confirmed data leakage— the inadvertent use of test-set information during training—in 23 (72%), collectively cited over 3,000 times [Asiaee et al., 2026]. Leakage artifacts include supervised feature screening performed before cross-validation, test-set early stopping, and pair-level splits inconsistent with cold-start generalization claims [Ambroise and McLachlan, 2002, Varma and Simon, 2006, Cawley and Talbot, 2010, Kapoor and Narayanan, 2023]. These evaluation errors systematically underestimate prediction error and inflate apparent model performance, making it difficult to assess whether proposed innovations yield genuine gains.

Independently, critical benchmarking studies have questioned the value proposition of increasingly complex architectures. Bernett et al. [2025] found that deep learning methods only modestly outperform a naive mean predictor under standardized evaluation. Branson et al. [2025] demonstrated that drug molecular structure contributes negligible predictive signal, with essentially all performance deriving from transcriptomic features. Li et al. [2023] showed that pathway-informed neural network architectures perform no better with biological pathways than with random gene groupings, challenging claims of built-in interpretability. Together, these findings suggest that the field’s emphasis on architectural novelty has outpaced the reliability of its evaluation practices.

A largely untapped source of legitimate inductive bias is tissue lineage. Cancer cell lines originate from distinct tissue types with different transcriptional programs, mutational landscapes, and drug vulnerabilities. Standard single-task models that pool all cell lines treat tissue heterogeneity as noise, potentially obscuring tissue-specific drug mechanisms while selecting features that reflect lineage confounding rather than drug-specific biology. The Tissue-guided LASSO (TG-LASSO) [Huang et al., 2020] incorporated tissue information by using non-target tissues as training data and the target tissue for model selection. However, TG-LASSO does not decompose the predictive signal into shared and tissue-specific components, limiting interpretability about which genomic features are universally versus selectively predictive.

Here we present the Data Shared Elastic Net (DSEN), a tissue-aware multitask regression that explicitly decomposes each drug’s regression coefficients into a shared component common to all tissue types and tissue-specific deviations [Strauch and Asiaee, 2024]. Building on the data-shared lasso framework [Gross and Tibshirani, 2016], DSEN adapts elastic-net regularization [Zou and Hastie, 2005, Friedman et al., 2010] for large-scale drug-by-drug modeling across tissue lineages. Crucially, all results are evaluated under leakage-free cross-validation with per-fold feature screening, ensuring that reported performance reflects genuine predictive ability rather than evaluation artifacts.

Our contributions are: (1) a tissue-aware decomposition that improves prediction for 92.5% of drugs under honest evaluation while producing sparser models; (2) head-to-head comparison showing that TG-LASSO, the only other tissue-aware method, performs substantially worse than even the tissue-agnostic baseline under leakage-free evaluation; (3) leave-tissue-out validation demonstrating that DSEN’s shared coefficients generalize to unseen tissue lineages; (4) feature modality ablation revealing that tissue-aware modeling provides the largest benefit when features are scarce, with no single modality dominating prediction; and (5) biologically coherent target recovery, with ubiquitous targets in the shared block and lineage-restricted targets in tissue-specific blocks.

## Methods

### Data sources

We analyzed drug sensitivity measurements from GDSC/Sanger release 6.0 and molecular features derived from CCLE/DepMap releases distributed with the CCLE update [Garnett et al., 2012, Iorio et al., 2016, Ghandi et al., 2019]. The predictor matrix contained 86,546 features for 1,462 cell lines representing 31 tissue lineages, spanning gene expression, copy number, mutation (four subtypes: hotspot, truncating, missense, hotspot-recurrent), and reverse-phase protein array (RPPA). For the RPPA modality, we retained all 231 antibody columns from the MD Anderson RPPA core; 139 of these are uniquely mapped to HGNC gene symbols and constitute the “RPPA” modality used throughout, while the remaining 92 antibodies—primarily those flagged by the MD Anderson QC pipeline (suffix `Caution` for known specificity concerns or `ValidationUnavailable` for antibodies lacking validation) or recognising protein isoforms without a one-to-one gene assignment (e.g. p14^ARF^, which shares the *CDKN2A* locus with p16^INK4a^)—are retained as predictors and labelled “Other RPPA” in figures. Drug response covered 265 compounds measured by the GDSC dose-response area under the curve (AUC), normalised to [0, 1], where larger values indicate greater resistance (less drug effect). Data provenance is documented in the accompanying code repository.

### Data Shared Elastic Net formulation

DSEN models drug response across *T* tissue types as related tasks. For each drug, we fit a single elastic-net model on a block-structured design that couples shared and tissue-specific coefficients. Specifically, for tissue *t* with *n*_*t*_ samples and *p* features, the coefficient vector is parameterized as *β*_*t*_ = *β*_shared_ + *β*_tissue,*t*_, where *β*_shared_ ∈ ℝ^*p*^ captures signal common to all tissues and *β*_tissue,*t*_ ∈ ℝ^*p*^ captures tissue-specific deviations. Equivalently, this is a model with feature main effects (the shared block) and tissue-by-feature interaction terms (the tissue-specific blocks), where the main and interaction effects are penalized differently according to the multiplier *ρ* introduced below: small *ρ* shrinks the interactions toward zero and recovers a pooled single-task fit, while large *ρ* frees the interactions to express lineage-specific structure. This shared-plus-deviation parameterization is the data-shared lasso construction of Gross and Tibshirani [2016], which we adapt here to elastic-net penalties and tissue-stratified, leakage-free evaluation; closely related shared/task-specific decompositions with theoretical guarantees have been analysed in the high-dimensional multitask setting [Asiaee et al., 2018, 2019].

To implement this decomposition within the standard glmnet framework [Friedman et al., 2010], we construct a sparse block-structured design matrix *Z* of dimension *N* × *p*(1 + *T* ), where *N* = ∑_*t*_ *n*_*t*_. For a sample from tissue *t*, the row of *Z* contains the original feature vector *x*_*i*_ in both the first *p* columns (shared block) and in columns *p* ·*t*+1 through *p* ·(*t*+1) (tissue-*t* block), with zeros elsewhere. The elastic-net objective becomes:

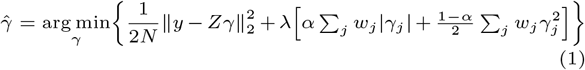

where 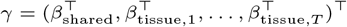 is the concatenated coefficient vector, *α* is the elastic-net mixing parameter, *λ* controls overall regularization strength, and *w*_*j*_ are feature-specific penalty weights.

The penalty weight vector *w* encodes the shared-versus-tissue tradeoff. Shared coefficients (first *p* entries) receive weight *ρ*, a hyperparameter controlling the relative penalization of shared structure. When *ρ >* 1, shared coefficients are penalized more heavily than tissue-specific ones, favoring tissue-specific structure; when *ρ <* 1, shared structure is favored. Tissue-specific coefficients are weighted proportionally 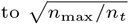, so that smaller tissues receive higher penalties, preventing overfitting in tissues with few samples.

Predictions for a new sample from tissue *t* are computed as 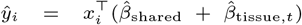. The decomposition is directly interpretable: 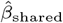 identifies features with pan-cancer predictive signal, while 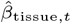 identifies features whose predictive role is specific to lineage *t*.

### Leakage-free cross-validation

All models were evaluated using 5-fold cross-validation with tissue-stratified folds. To prevent data leakage, the complete preprocessing pipeline was repeated independently within each fold using training data only [Asiaee et al., 2026, Ambroise and McLachlan, 2002]:

1. **Variance filtering**: retain features with variance *>* 0.01 (computed on training samples).
2. **Correlation screening**: retain features with |cor(*x*_*j*_, *y*)| *>* 0.1 (computed on training samples).
3. **Duplicate removal**: remove duplicated columns after filtering.
4. **Scaling**: z-score continuous features using training-set mean and standard deviation; binary mutation features were excluded from scaling.

The same transformations were applied to held-out test samples using training-set statistics. Tissues with fewer than 25 cell lines per drug were excluded from DSEN modeling to ensure stable within-tissue estimation.

Hyperparameters were selected by grid search over the elastic-net mixing parameter *α* ∈ {0.05, 0.1, …, 1.0} (12 values) and the shared-block penalty multiplier *ρ* ∈ {0.125, 0.25, 0.5, 1, 2, 4}. The combination minimizing 5-fold CV mean squared error was selected. For the standard elastic-net baseline, the same *α* grid and leakage-free protocol were used without the tissue decomposition.

Performance is reported as MSE on held-out folds. Throughout, MSE improvement is defined as 100 × (MSE_EN_ − MSE_method_)/MSE_EN_, so that positive values denote lower error than the standard elastic-net baseline and negative values denote higher error.

### Bootstrap feature selection

To characterize feature selection behavior, we performed bootstrap resampling (200 replicates) and recorded how often each feature received a non-zero coefficient. A feature was considered “stable” if selected in at least 50% of bootstrap resamples (unless otherwise noted). For DSEN, we tracked shared and tissue-specific coefficients separately: a feature was tissue-selected if its bootstrap frequency exceeded the threshold in any tissue. Combined DSEN selection was defined as the union of shared and tissue-specific stable feature sets.

### Drug–target curation and target recovery

Drug–target relationships were curated from GDSC metadata. Gene names were standardized using a curated alias-to-HGNC mapping. Of 265 drugs, 220 had at least one gene/protein target mappable to the feature space; the remaining 45 drugs were excluded from target-recovery analyses (3 lacked any GDSC target annotation and the rest had only non-gene targets such as “microtubule stabilizer”).

Feature-to-gene mapping accounted for all modalities: gene expression, copy number, mutation subtypes, and 139 RPPA features using antibody-to-gene mappings (e.g., B-Raf pS445 → *BRAF*, Akt pS473 → *AKT1* ). A complete RPPA-to-HGNC mapping is provided in the code repository.

Target recovery was evaluated by checking whether a drug’s known target gene appeared among its stable selected features. For DSEN, shared-block and tissue-specific recovery were assessed separately, and combined recovery was defined as the union.

### Biological validation

To characterise the lineage breadth of selected features, each stably-selected gene was assigned the tissue-level expression-specificity score 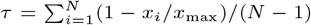, where *x*_*i*_ is the gene’s mean log-expression in tissue *i, x*_max_ is its maximum tissue-level mean across the *N* = 31 GDSC lineages, and lower values of *τ* indicate broader expression across lineages while higher values indicate stronger lineage restriction. Differences in median *τ* between method blocks were tested with a two-sided Wilcoxon rank-sum test. Aggregate per-block breadth statistics are reported in Supplementary Table S1.

### Comparison methods

#### Standard elastic net

A single elastic-net model fitted to all cell lines (ignoring tissue labels), using the same leakage-free CV protocol and hyperparameter grid described above.

#### TG-LASSO

Tissue-guided LASSO [Huang et al., 2020] is the most directly comparable tissue-aware method. For each tissue in the training fold, TG-LASSO trains a LASSO model on all *non-target* tissues and selects *λ* by minimizing MSE on the target tissue (used as a validation set). At test time, samples from each tissue are predicted by the model trained for that tissue. We implemented TG-LASSO using the same leakage-free CV protocol (per-fold feature screening, same tissue-stratified folds) and evaluated it on all 265 drugs. Key methodological differences from DSEN: TG-LASSO excludes target-tissue samples from model fitting (using them only for hyperparameter selection), whereas DSEN jointly trains on all tissues with shared and tissue-specific coefficients; TG-LASSO uses pure LASSO (*α* = 1), whereas DSEN uses elastic-net regularization.

## Results

### DSEN improves prediction under leakage-free evaluation

Removing data leakage from the evaluation pipeline provides an honest baseline against which genuine modeling innovations can be tested. We compared DSEN to a standard elastic net across 265 drugs under leakage-free cross-validation with tissue-stratified folds.

DSEN improved MSE relative to the standard elastic net for 92.5% of compounds (245/265), with a mean improvement of 4.95% (median 3.95%; Fig. 1a; Table 1). These gains are measured against the leakage-free baseline, confirming that tissue-aware structure provides genuine predictive benefit rather than recovery of leaked optimism. The maximum improvement reached 22.7% (GNF-2), and only 20 drugs (7.5%) showed higher MSE under DSEN.

**Table 1.** DSEN performance summary under leakage-free evaluation. Key metrics comparing standard elastic net and DSEN across 265 drugs with tissue-stratified 5-fold CV. Win rate indicates the fraction of drugs where DSEN achieves lower MSE. Feature counts use a 50% bootstrap frequency threshold (200 resamples). Target recovery is evaluated over 220 drugs with at least one mapped gene target.

| Metric | Value |
| --- | --- |
| *MSE performance* |  |
| Win rate | 92.5% |
| Mean improvement | 4.95% |
| Median improvement | 3.95% |
| Max improvement (GNF-2) | 22.7% |
| *Model structure* |  |
| Drugs with $\rho < 1$ (shared-favoring) | 17.4% |
| Drugs with $\rho \geq 1$ (tissue-favoring) | 82.6% |
| Drugs with zero shared features | 23.4% |
| DSEN shared sparser than EN | 63.8% |
| Mean stable features: EN | 29.0 |
| Mean stable features: DSEN shared | 12.2 |
| Mean stable features: DSEN tissue | 33.7 |
| *Target recovery (220 drugs)* |  |
| Any target (0.005 threshold): EN / DSEN combined | 50.5% / 63.6% |
| Any target (0.5 threshold): EN / DSEN combined | 8.6% / 10.5% |
| Drugs with tissue-only recovery | 6 |

**Fig 1.**
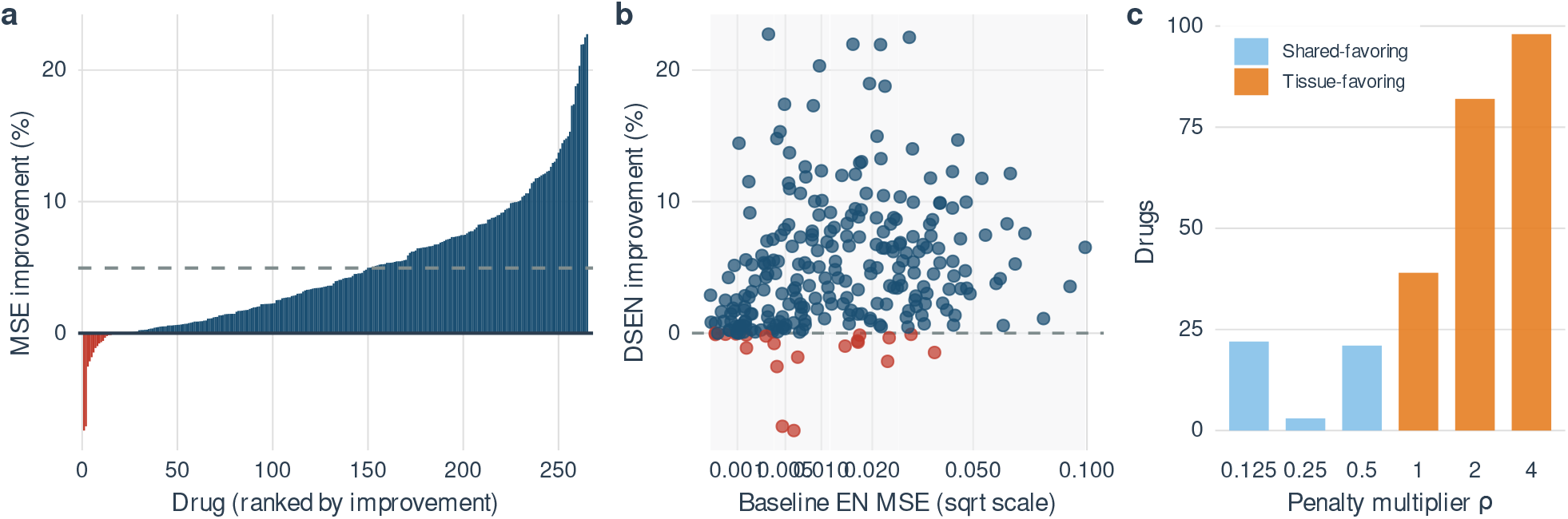
DSEN improves leakage-free drug response prediction across 265 drugs. **(a)** Waterfall plot of per-drug MSE improvement (%) for DSEN relative to a standard elastic net under leakage-free CV; blue bars indicate DSEN wins (92.5%, 245/265 drugs) and red bars indicate EN wins (7.5%, 20/265); the dashed line marks the mean improvement (4.95%). **(b)** Improvement versus baseline elastic-net MSE; light bands denote difficulty quartiles (Q1 89.6%, Q2 92.4%, Q3 90.9%, Q4 97.0% win rate). Points are colored by winner. **(c)** Distribution of the shared-block penalty multiplier *ρ*; values *<* 1 (blue) favor shared structure, values ≥ 1 (orange) favor tissue-specific coefficients. The majority of drugs select *ρ* ≥ 1. *Alt text:* Three panels: (a) a waterfall bar plot of per-drug MSE improvement, predominantly positive; (b) a scatter of DSEN improvement against baseline elastic-net MSE, points coloured by winning method; (c) a bar chart of the selected shared-block penalty multiplier *ρ*, with most drugs at *ρ* ≥ 1.

When drugs were stratified by baseline prediction difficulty (elastic-net MSE quartile), DSEN showed consistent improvement across all ranges: Q1 (easiest) 89.6% win rate with 2.65% mean gain; Q2 92.4% with 5.08%; Q3 90.9% with 6.39%; and Q4 (hardest) 97.0% with 5.73% (Fig. 1b). The high win rate in the hardest quartile suggests that tissue-specific structure is particularly valuable when pan-cancer signal alone is insufficient for accurate prediction.

The selected shared-block penalty multiplier *ρ* revealed the balance between shared and tissue-specific structure (Fig. 1c). The majority of drugs (82.6%) selected *ρ* ≥ 1, favoring tissue-specific coefficients, and 68% selected *ρ* ≥ 2. Only 17.4% of drugs favored shared structure (*ρ* < 1). This distribution indicates that tissue-specific signal carries substantial predictive value for most compounds, consistent with the biological expectation that drug mechanisms vary across lineages.

### DSEN produces sparser shared models

Beyond prediction accuracy, DSEN reorganizes model structure by redistributing predictive features between shared and tissue-specific blocks. In bootstrap analyses (features selected in ≥50% of 200 resamples), DSEN’s shared coefficient block was sparser than the standard elastic net for 63.8% of drugs (Fig. 2b). The shared block contained a mean of 12.2 stable features versus 29.0 for the standard elastic net, a 58% reduction.

**Fig 2.**
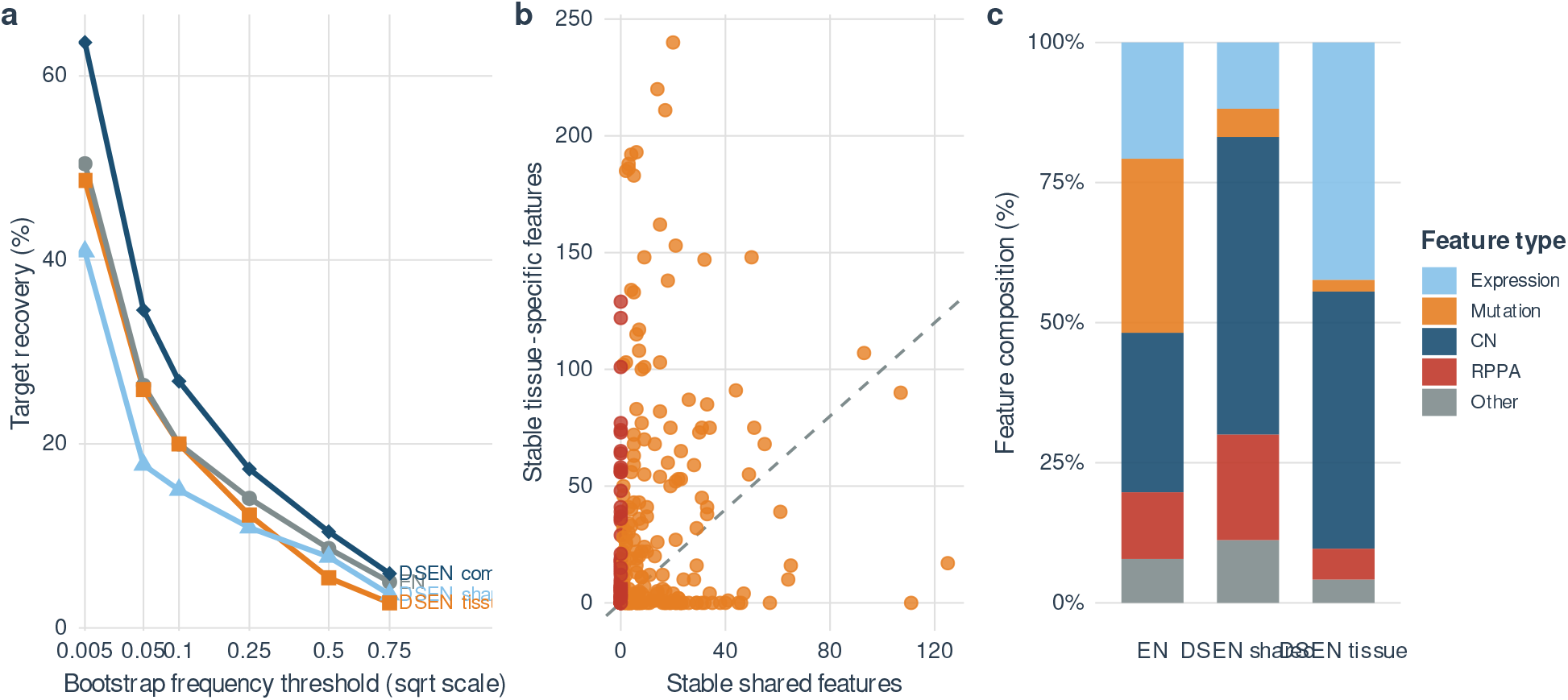
DSEN model structure, target recovery, and feature composition. **(a)** Target recovery rates across bootstrap frequency thresholds for EN (gray), DSEN shared (light blue), DSEN tissue (orange), and DSEN combined (dark blue). DSEN combined exceeds EN at all thresholds. **(b)** Stable shared versus tissue-specific feature counts per drug at the 0.5 threshold; orange: drugs with non-zero shared features; red: drugs with zero shared features (23.4%). The shared block is sparser than EN for 63.8% of drugs. **(c)** Feature modality composition of stable features for EN, DSEN shared, and DSEN tissue-specific blocks. “Other” denotes RPPA antibody features not mapped to an HGNC gene in the curated 139-antibody panel (92 antibodies). *Alt text:* Three panels: (a) target-recovery curves versus bootstrap-frequency threshold for EN, DSEN shared, DSEN tissue and DSEN combined, with DSEN combined highest; (b) a scatter of stable shared versus tissue-specific feature counts per drug; (c) stacked bars of feature-modality composition for EN, DSEN shared and DSEN tissue blocks.

DSEN additionally assigns tissue-specific deviations (mean 33.7 stable tissue-specific features), but these are localized to individual lineages rather than contributing to a universal signature. For 23.4% of drugs (62/265), the shared block contained zero stable features, indicating predominantly tissue-specific predictive structure (Fig. 2b). These drugs may have mechanisms of action that are entirely lineage-dependent, or they may require tissue context to resolve predictive features from background noise.

The sparsity of the shared block has practical implications: a smaller universal feature set is easier to interpret biologically and more amenable to translation into clinical assays. Rather than a monolithic model selecting dozens of features of uncertain generalizability, DSEN partitions the signal so that shared features can be interpreted as candidate pan-cancer markers while tissue-specific features provide lineage-resolved mechanistic hypotheses.

### Improved target recovery through tissue decomposition

A key test of biological coherence is whether selected features recover known drug targets. We evaluated target recovery across 220 drugs with at least one mapped gene target in the feature space, using bootstrap frequency thresholds ranging from 0.005 (most permissive) to 0.75 (most stringent).

Under strong regularization and a stringent stability threshold (0.5), direct target recovery was rare for both methods, consistent with polygenic drug sensitivity and heavy penalization. The standard elastic net recovered any known target for 8.6% of drugs (19/220), while DSEN’s shared block recovered targets for 7.7% (17/220). However, when tissue-specific coefficients were also considered, combined DSEN recovery rose to 10.5% (23/220; Fig. 2a; Table 1); selected example recoveries are listed in Table 3. The tissue block contributed a small but real set of additional target recoveries, including primary curated targets such as Rapamycin–MTOR and Sunitinib–PDGFRA, as well as lineage-resolved secondary targets such as Ruxolitinib–JAK2 and PD173074–FGFR2. Thus, the main biological value of the tissue block is not a large aggregate jump in direct target recall, but the ability to assign selected genes to lineage-specific contexts.

DSEN combined recovery exceeded the elastic net at every threshold tested, from 63.6% versus 50.5% at the most permissive threshold (0.005) to 5.9% versus 5.0% at the most stringent (0.75; Fig. 2a). The consistent advantage across thresholds indicates that DSEN’s tissue decomposition captures additional target-associated signal in tissue-specific coefficients that a single-task model cannot access.

### Systematic biological validation of the decomposition

To move beyond isolated target examples, we mapped stable features (≥ 50% of 200 bootstrap resamples) to genes and compared lineage breadth using tissue-median expression across the same CCLE/DepMap cell lines used for model fitting. Shared-block selections were modestly but significantly broader than tissue-block selections (median expression-specificity score *τ* = 0.136 versus 0.147; Wilcoxon *p* = 0.040), with shared genes more often falling in the broadest quartile of lineage breadth (15.9% versus 10.7%; full per-block breadth statistics in Supplementary Table S1). This shift was real but not absolute, indicating that DSEN does not create a clean pan-cancer versus lineage-restricted partition for every recovered feature.

The stronger biological signal emerged at the level of recurrent pathway and lineage programs (Fig. 3; Fig. 4). Nutlin-3a recovered a four-gene shared *MDM2* /*TP53* /*BAX* /*CDKN1A* module, matching canonical p53-pathway activation by MDM2 antagonism [Vassilev et al., 2004]. The signs of the shared coefficients are biologically coherent (Fig. 4a): the *TP53* - mutation feature carries a large positive coefficient, capturing the well-established resistance of *TP53* -mutant cells to MDM2 inhibition (mutant p53 cannot be reactivated, so apoptosis fails), while *MDM2, CDKN1A* and *BAX* protein levels carry negative coefficients—consistent with cells that have abundant drug target and a primed p53 transcriptional output being more sensitive.

**Fig 3.**
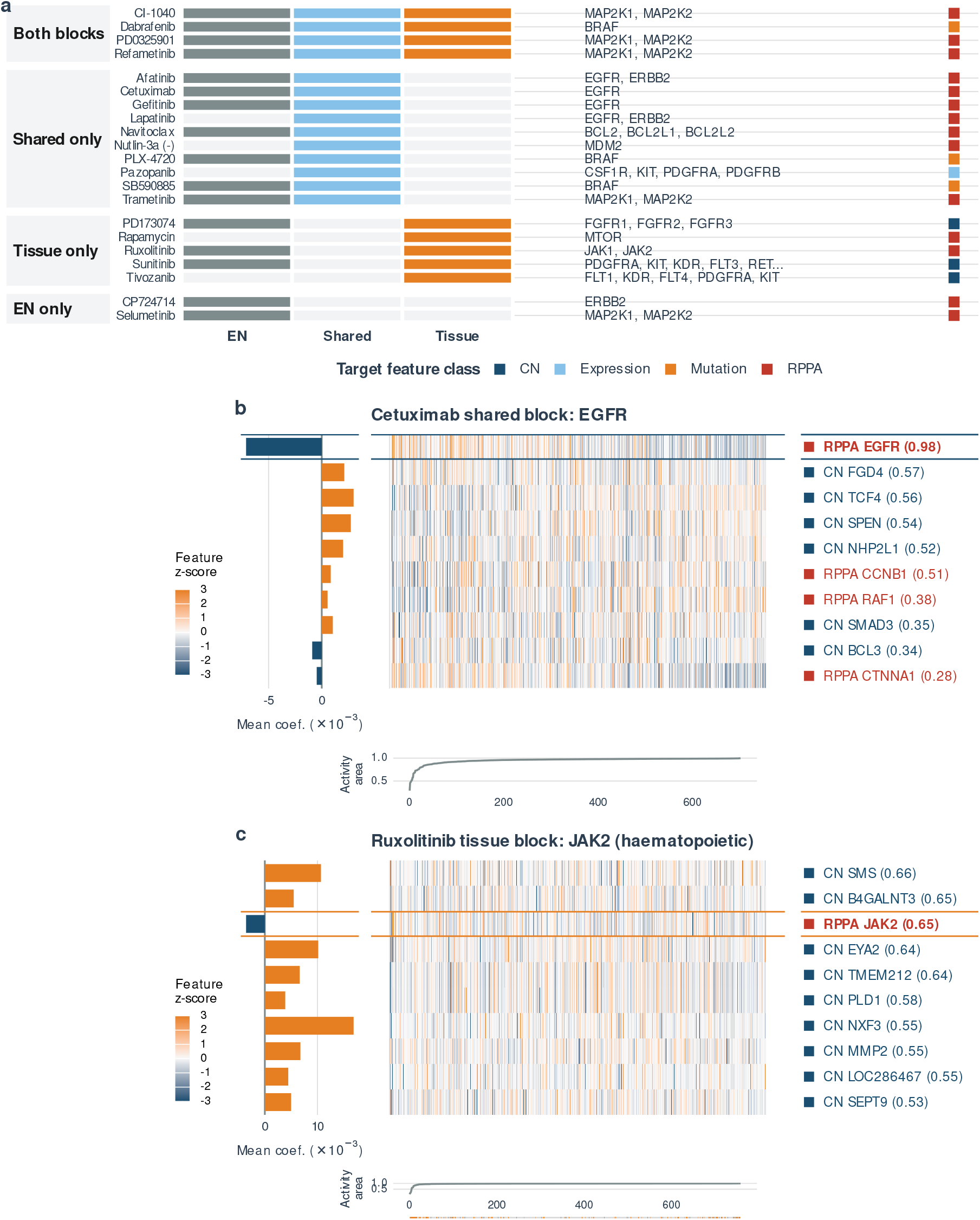
Target recovery decomposition for selected drugs. **(a)** Overview matrix for drugs with recovered targets at the 0.5 threshold, grouped by recovery location (both blocks, shared only, tissue-specific only, EN only). **(b)** Shared-block exemplar: Cetuximab with *EGFR* recovered as a shared feature. Left: signed shared coefficients; center: z-scored feature values across cell lines ordered by drug response; right: bootstrap selection frequencies. The *EGFR* protein coefficient is negative, so cells with higher EGFR protein are predicted to have lower AUC, i.e. to be more sensitive to EGFR blockade. **(c)** Tissue-specific exemplar: Ruxolitinib with *JAK2* recovered only in the haematopoietic/lymphoid tissue block, absent from the shared block—consistent with the lineage-restricted role of activating *JAK2* lesions in myeloproliferative disease. *Alt text:* Three panels: (a) an overview matrix grouping drugs by where their known targets were recovered (both blocks, shared only, tissue only, EN only); (b) Cetuximab, showing EGFR recovered as a shared feature with its coefficient, feature values and bootstrap frequency; (c) Ruxolitinib, showing JAK2 recovered only in the haematopoietic/lymphoid tissue block.

**Fig 4.**
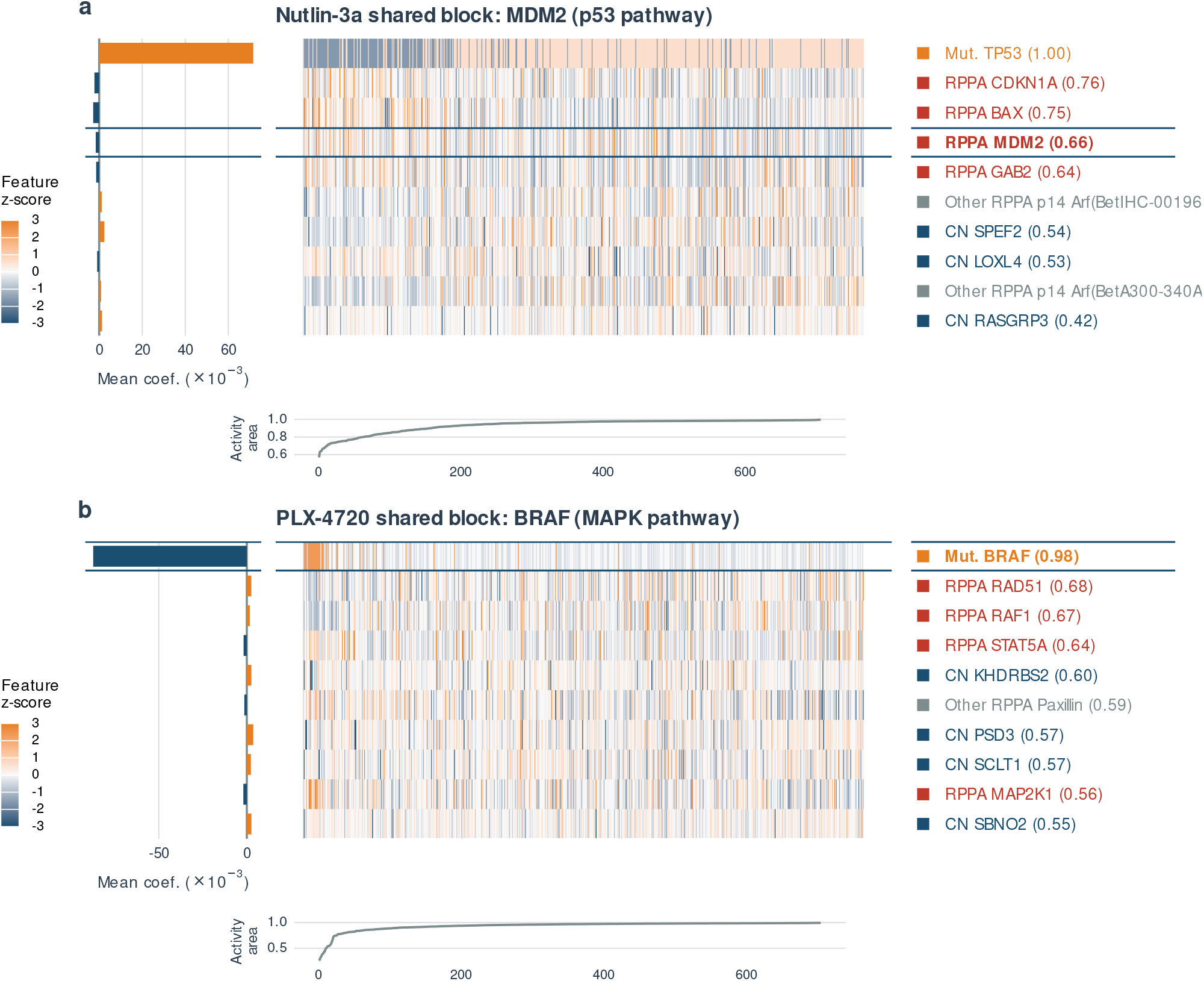
Shared-block exemplars for recurrent pathway modules. **(a)** Nutlin-3a, with *MDM2* recovered as a shared feature (p53 pathway). Left: signed shared coefficients; centre: z-scored feature values across cell lines ordered by drug response; right: bootstrap selection frequencies. The *TP53* - mutation feature is strongly positive (mutant p53 cannot be reactivated by MDM2 inhibition, raising AUC and conferring resistance), while *MDM2, CDKN1A* and *BAX* protein levels are negative (more drug target and a primed p53 output predict lower AUC, i.e. greater sensitivity). **(b)** PLX-4720, with *BRAF* recovered as a shared feature (MAPK pathway), shown in the same layout. The *BRAF* -mutation coefficient is strongly negative, capturing the canonical sensitivity of V600E-mutant cells to BRAF inhibition; *MAP2K1* (MEK1) is mildly negative, reflecting MAPK-pathway dependence; and *RAF1* (CRAF) is positive, consistent with paradoxical RAF1-mediated reactivation of MAPK signalling that drives resistance in BRAF-wildtype cells. Larger AUC values indicate greater resistance throughout. *Alt text:* Two stacked panels showing shared-block exemplars for Nutlin-3a (panel a, with MDM2/TP53/CDKN1A/BAX features) and PLX-4720 (panel b, with BRAF/RAF1/MAP2K1 features); each panel has a coefficient bar plot on the left, a z-scored heatmap of feature values across cell lines in the centre, and feature labels with bootstrap frequencies on the right.

PLX-4720 similarly recovered a shared MAPK module consisting of *BRAF, RAF1*, and *MAP2K1* ; *BRAF* mutations are recurrent oncogenic drivers across multiple cancer types [Davies et al., 2002], and selective BRAF inhibition was originally developed for melanoma while retaining pathway specificity [Tsai et al., 2008]. Coefficient signs again align with the canonical biology (Fig. 4b): the *BRAF*-mutation feature is strongly negative, capturing the exquisite sensitivity of V600E-mutant cells to BRAF inhibition; *MAP2K1* (MEK1) protein is mildly negative, indicating MAPK-pathway dependence; and *RAF1* (CRAF) protein has a positive coefficient, consistent with the well-documented paradoxical activation of MAPK signalling by ATP-competitive BRAF inhibitors in cells with active CRAF, which contributes to resistance in BRAF-wildtype backgrounds. More generally, *MAP2K1* appeared in the shared block for five MAPK-pathway drugs, supporting interpretation of the shared block as a transferable pathway component rather than only direct-target capture.

The clearest lineage-specific pattern was observed for MAPK-pathway drugs in skin. Across 11 BRAF/MEK-pathway compounds and replicates, the skin tissue block repeatedly recovered melanoma-lineage markers including *MITF* (8 drugs), *S100B* (4), and *PLP1* (4), alongside skin-specific *BRAF* recovery in three models. *MITF* is a melanoma lineage-survival oncogene [Garraway et al., 2005], and *S100B* is a well-established melanoma marker [Nonaka et al., 2008]. Consistent with this lineage dependence, leave-skin-out performance for the same MAPK set was negative in 10 of 11 cases (median −19.2% for DSEN shared versus EN; Supplementary Table S2), linking the tissue block to a concrete generalization failure mode rather than to anecdotal target recovery.

Among single-drug tissue-specific target examples, *JAK2* was recovered only in the haematopoietic/lymphoid block for Ruxolitinib, consistent with the lineage-restricted role of activating *JAK2* lesions in myeloproliferative disease [James et al., 2005]. Rapamycin similarly recovered *MTOR* only in the haematopoietic/lymphoid block. Some candidate examples were not stable at the primary threshold: PD173074 recovered *FGFR2* rather than *FGFR3*, and the main AKT inhibitor VIII model did not stably recover *AKT1*. Taken together, these results support a cautious interpretation: DSEN separates transferable pathway signal from lineage-specific modifiers, but the strongest evidence lies in recurrent pathway and tissue programs rather than in a universal gene-by-gene partition.

### Feature modality analysis

Analysis of stable feature composition revealed distinct modality profiles across model components (Fig. 2c). The standard elastic net and DSEN shared block were dominated by gene expression and copy number features, consistent with these modalities providing the broadest genomic coverage. DSEN tissue-specific blocks showed a relative enrichment for RPPA (protein) features compared to the shared block. However, the feature-modality ablation below revealed that this enrichment has limited predictive consequence: removing all 139 RPPA features increased MSE by only 0.35%, and expression of the same genes outperformed RPPA measurements for the majority of drugs.

### Comparison with TG-LASSO

Under leakage-free evaluation on the same 265 drugs and tissue-stratified folds, TG-LASSO performed substantially worse than both the standard elastic net and DSEN (Table 2). TG-LASSO achieved lower MSE than EN for only 1 of 265 drugs (CP724714, +3.4%), with a mean MSE *increase* of 13.8% relative to EN (median 9.8%). DSEN outperformed TG-LASSO on all 265 drugs.

**Table 2.** Method comparison under leakage-free evaluation. Comparison of standard elastic net, TG-LASSO, and DSEN across 265 drugs. All methods evaluated with identical leakage-free CV protocol and tissue-stratified folds. TG-LASSO performed substantially worse than even the tissue-agnostic EN baseline, winning on only 1 of 265 drugs.

| Metric | EN | TG-LASSO | DSEN |
| --- | --- | --- | --- |
| Mean MSE | 0.01716 | 0.02004 | **0.01616** |
| Win rate vs EN (%) | — | 0.4 | **92.5** |
| Mean MSE improvement vs EN (%) | — | −13.8 | **+4.95** |
| Median MSE improvement vs EN (%) | — | −9.8 | **+3.95** |
| Interpretable decomposition | No | No | Yes |

**Table 3.**
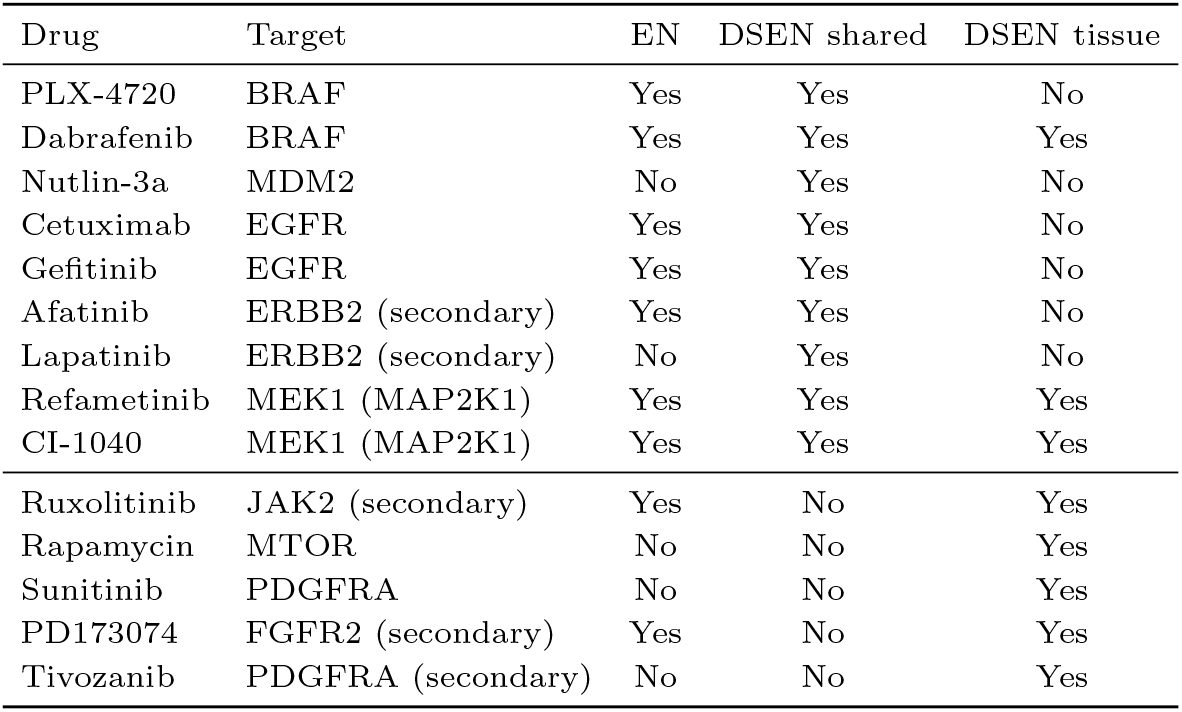
Selected direct target recoveries at the 50% bootstrap threshold. Features selected in ≥50% of 200 bootstrap resamples. “Secondary” denotes a curated secondary target from the GDSC-derived target table.

| Drug | Target | EN | DSEN shared | DSEN tissue |
| --- | --- | --- | --- | --- |
| PLX-4720 | BRAF | Yes | Yes | No |
| Dabrafenib | BRAF | Yes | Yes | Yes |
| Nutlin-3a | MDM2 | No | Yes | No |
| Cetuximab | EGFR | Yes | Yes | No |
| Gefitinib | EGFR | Yes | Yes | No |
| Afatinib | ERBB2 (secondary) | Yes | Yes | No |
| Lapatinib | ERBB2 (secondary) | No | Yes | No |
| Refametinib | MEK1 (MAP2K1) | Yes | Yes | Yes |
| CI-1040 | MEK1 (MAP2K1) | Yes | Yes | Yes |
| Ruxolitinib | JAK2 (secondary) | Yes | No | Yes |
| Rapamycin | MTOR | No | No | Yes |
| Sunitinib | PDGFRA | No | No | Yes |
| PD173074 | FGFR2 (secondary) | Yes | No | Yes |
| Tivozanib | PDGFRA (secondary) | No | No | Yes |

The performance gap reflects a fundamental design difference: TG-LASSO excludes target-tissue samples from model fitting, using them only for *λ* selection. While this prevents tissue-specific overfitting, it also discards the within-tissue predictive signal that both EN and DSEN retain. Under leakage-free evaluation, this conservative strategy proves counterproductive—the validation set for each tissue (25–100 samples from a single tissue within one training fold) provides a noisy estimate of *λ*, while the loss of training data degrades model quality.

TG-LASSO was originally evaluated with pair-level cross-validation [Huang et al., 2020], which has been identified as a form of data leakage for cold-start prediction [Asiaee et al., 2026]. Under honest evaluation, TG-LASSO’s tissue-guided approach does not improve upon simply pooling all tissues, while DSEN’s joint decomposition into shared and tissue-specific components captures tissue structure without sacrificing training data.

### Leave-tissue-out generalization

To test whether DSEN’s shared coefficients capture genuinely pan-cancer signal, we performed leave-tissue-out (LTO) validation: for each drug, we held out one tissue entirely, trained DSEN on the remaining tissues, and predicted the held-out tissue using only the shared coefficient block 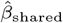 (the held-out tissue has no tissue-specific block since it was absent from training). This was evaluated across 224 drugs with ≥ 5 tissues, yielding 2,800 tissue-holdout predictions.

DSEN’s shared block achieved lower MSE than the tissue-agnostic EN for 58.8% of tissue-holdout predictions, with a mean per-drug MSE improvement of 1.8%. At the drug level, 149 of 224 drugs (66.5%) had DSEN winning on more than half their tissues. Per-tissue results varied (Supplementary Table S3): DSEN’s shared block showed strongest generalization to large intestine (+17.1%), upper aerodigestive tract (+6.5%) and central nervous system (+4.7%), and slightly worse generalization to bone (−2.2%) and oesophagus (−1.4%). Averaged across all 211 drugs evaluable on skin, leave-skin-out generalization was also positive (+4.9%); the aggregate, however, conceals a striking subset failure for MAPK-pathway drugs, discussed below.

The modest but consistent LTO advantage confirms that DSEN’s shared block captures signal that transfers to unseen tissues—a stronger test of generalization than standard cross-validation. The drug-class-specific losses are also biologically informative. In particular, although the aggregate leave-skin-out result was positive, skin was a recurrent failure mode within the MAPK-pathway subset: across the 11 strict BRAF/MEK compounds and replicates, leave-skin-out performance was negative in 10 of 11 cases (median −19.2%; Supplementary Table S2), matching the repeated recovery of skin-specific *MITF* /*S100B* /*PLP1* tissue blocks described above. This suggests that the shared block captures the transferable part of MAPK dependence, while the tissue block absorbs melanoma-lineage modulation that does not generalize when skin is withheld.

### Feature modality ablation

To assess the contribution of each feature modality, we ran the full DSEN pipeline on 12 feature subsets using identical folds and evaluation protocol (Table 4). DSEN consistently improved over EN regardless of feature composition, with win rates ranging from 84.6% (517 RPPA-gene multi-modal features) to 94.0% (112 RPPA-gene expression features).

**Table 4.** Feature modality ablation. DSEN performance under different feature subsets, all evaluated with identical folds and leakage-free CV. Win rate and improvement are DSEN vs standard EN within each feature subset. Tissue decomposition benefits prediction across all modalities, with the largest relative improvement when features are scarce.

| Feature subset | Features | N drugs | Win rate (%) | Mean impr. (%) |
| --- | --- | --- | --- | --- |
| All features | 86,546 | 265 | 92.5 | 4.95 |
| No RPPA | 86,407 | 265 | 91.3 | 4.92 |
| No phospho-RPPA | 86,507 | 265 | 92.1 | 4.97 |
| No total-RPPA | 86,446 | 265 | 92.5 | 4.93 |
| Expr + Mut + CN | 86,315 | 265 | 90.6 | 4.71 |
| Expression only | 18,960 | 248 | 92.7 | 3.57 |
| Mutation only | 44,039 | 265 | 92.8 | 5.53 |
| RPPA only | 139 | 261 | 91.2 | 3.05 |
| RPPA phospho only | 39 | 219 | 89.5 | 2.12 |
| RPPA total only | 100 | 256 | 91.4 | 3.44 |
| RPPA-gene multi-modal | 517 | 260 | 84.6 | 3.60 |
| RPPA-gene expression | 112 | 167 | **94.0** | **6.25** |

Removing all 139 RPPA features from the full model increased DSEN MSE by only 0.35%. Removing only phospho-protein features (39 antibodies) had negligible effect (+0.05%). RPPA features alone (139 features) achieved a DSEN win rate of 91.2%, demonstrating that tissue structure benefits prediction even with a small, protein-only feature set. However, the absolute MSE with RPPA-only features was 5.6% higher than with all features.

On the 166 drugs where both RPPA protein features and expression of the same 112 genes could be evaluated, gene expression outperformed RPPA protein measurements on 106 drugs (63.9%) in terms of absolute DSEN MSE. This suggests that for these genes, mRNA levels provide comparable or better predictive information than protein abundance or phosphorylation state, likely due to expression measurements capturing a broader dynamic range of variation across cell lines.

The most striking ablation finding is that tissue-aware modeling provides the *largest* relative improvement when features are scarce. DSEN with only the 112 expression features corresponding to RPPA-panel genes achieved a 6.25% mean improvement over EN—the highest improvement rate of any subset. This indicates that DSEN’s tissue decomposition is most valuable precisely when the feature space is limited and tissue-specific signal would otherwise be obscured by noise.

## Discussion

We have presented DSEN, a tissue-aware elastic-net regression that decomposes drug response prediction into shared and tissue-specific components. Evaluated under leakage-free cross-validation—a prerequisite that 72% of published methods fail to satisfy [Asiaee et al., 2026]—DSEN improves prediction for the vast majority of drugs while reorganizing model structure in biologically interpretable ways.

### Interpretability as a feature, not a limitation

In a field where deep learning models have been shown to barely exceed mean-predictor baselines under honest evaluation [Bernett et al., 2025] and where pathway-informed architectures offer no advantage over random gene groupings [Li et al., 2023], DSEN’s transparent coefficient decomposition is a substantive advantage rather than a concession. The biological validation presented here supports a careful interpretation of that decomposition. Aggregate breadth comparisons showed only a modest shift between shared and tissue blocks, so the data do not justify claiming that every shared feature is broadly pan-cancer or that every tissue feature is sharply lineage-restricted. What is supported is more specific and more useful: shared coefficients recurrently recover transferable pathway modules such as the p53 and MAPK axes, whereas tissue coefficients repeatedly recover lineage markers that explain where shared structure fails to transfer, especially the melanoma-associated *MITF* /*S100B* program in skin. This coefficient-level decomposition provides a more direct mechanistic readout than black-box predictors alone.

### Clinical relevance of feature modalities

DSEN can operate on molecular feature classes that overlap with translational profiling: somatic mutations, copy number alterations, expression panels, and protein measurements where available. Feature ablation revealed that no single modality dominates: removing RPPA protein features increased MSE by only 0.35%, and gene expression of the same 112 genes outperformed RPPA protein measurements on 64% of drugs evaluated head-to-head. This suggests that the field’s overwhelming reliance on gene expression [Branson et al., 2025] is not unreasonable for prediction accuracy, though protein-level features may still offer complementary value for biological interpretation. More importantly, DSEN’s tissue decomposition provided the largest relative improvement when features were scarce (6.25% with only 112 features), suggesting that modeling tissue context may be more impactful than adding feature modalities.

### Resistance and protection signals

An exploratory observation from the coefficient analysis is that some drugs appear to recover a sign-imbalanced set of stable features, with more features predicting higher AUC (a proxy for resistance, in the convention used here) than lower AUC. We have not characterised this asymmetry quantitatively in the present work, and the apparent imbalance could partly reflect feature–feature correlation structure rather than mechanism; we therefore offer it only as a direction for future analysis. If it survives further scrutiny, the signed coefficient blocks could inform combination therapy design by surfacing features whose presence predicts resistance rather than response.

### The tissue definition challenge

DSEN requires discrete tissue labels, which are straightforward for established cell line panels (CCLE, GDSC) but less clear for patient cohorts or newer screening platforms such as PRISM. Extending DSEN to molecular subtypes (e.g., basal vs. luminal breast cancer) or to data-driven cell line groupings could capture finer-grained heterogeneity. Adaptive approaches that learn the grouping jointly with the model represent a natural extension.

### Relationship to TG-LASSO

TG-LASSO [Huang et al., 2020] is the most directly comparable tissue-aware method. Both leverage tissue information, but differ fundamentally in strategy. TG-LASSO uses tissue as a data-splitting device: it trains on non-target tissues and uses the target tissue solely for hyperparameter selection, effectively discarding within-tissue training signal. Under leakage-free evaluation, this approach performed substantially worse than even the tissue-agnostic elastic net (−13.8% mean MSE, winning on only 1 of 265 drugs). DSEN’s joint modeling—retaining all tissues for training while decomposing coefficients into shared and tissue-specific blocks—preserves training data while still capturing tissue-specific structure. The contrast illustrates that how tissue information is incorporated matters as much as whether it is used at all: tissue-aware modeling can harm prediction if it reduces effective sample size, while DSEN’s additive decomposition avoids this tradeoff.

### Limitations

DSEN belongs to the elastic-net family and is inherently linear, which may limit its ability to capture nonlinear drug–gene interactions. However, given that complex nonlinear models have not demonstrated convincing advantages under honest evaluation [Bernett et al., 2025, Branson et al., 2025], linearity may be a reasonable modeling choice for the current data regime. The tissue decomposition depends on available annotations and cannot capture subtissue heterogeneity without additional labels. Target recovery, while improved by tissue decomposition, remains low overall (∼10% at the 0.5 threshold), reflecting the polygenic nature of drug response and the limitations of penalized regression for recovering individual causal features from high-dimensional, correlated feature spaces. Our companion audit [Asiaee et al., 2026] further showed that pre-CV feature screening and pair-level splits inflate not only predictive accuracy but also reported target-recovery rates; the numbers reported here are therefore deliberately conservative relative to those obtained under leaky evaluation in the prior literature.

### Future directions

Natural extensions include adaptive sharing across lineages or molecular subtypes, sharing information across related drugs (for example compounds with shared targets or mechanisms of action) so that signal is borrowed across compounds as well as across tissues, integration with patient-derived xenograft or clinical trial data where tissue labels are available, and combination with stability selection frameworks that formally control false discovery. The DSEN decomposition could also serve as an interpretability layer for more complex models: shared and tissue-specific coefficient blocks can provide biological priors or initialization for tissue-aware neural networks.

## Conclusion

In a field where the majority of published methods have confirmed evaluation artifacts, DSEN provides a transparently evaluated, biologically interpretable approach to tissue-aware drug response modeling. By decomposing the predictive signal into shared and tissue-specific components, DSEN improves prediction for 92.5% of drugs under leakage-free assessment while producing sparser, more interpretable models. The most directly comparable tissue-aware method (TG-LASSO) performs substantially worse than even the tissue-agnostic baseline under leakage-free evaluation, while DSEN’s shared coefficients generalize to unseen tissues. Feature ablation confirms that tissue-aware modeling—not any particular feature modality—is the primary driver of DSEN’s improvement, with the benefit increasing as the feature space shrinks. The strongest biological evidence for the decomposition is recurrent pathway recovery in shared blocks together with recurrent lineage markers in tissue blocks, especially the skin-specific MAPK program captured by *MITF, S100B*, and *PLP1*. DSEN’s transparent coefficient structure positions it as a practical tool for biomarker nomination and assay design in precision oncology.

## Acknowledgements

We thank Mahmoud Ghandi for sharing original CCLE analysis code and data.

## Author contributions

J.S. developed and implemented DSEN and drafted the first version of the manuscript. L.A. drafted portions of the manuscript. H.H.P. provided biological interpretation and edited the manuscript. J.P.L. contributed statistical methodology guidance and edited the manuscript. K.R.C. supervised early stages of the project and edited the manuscript. A.A. conceived the study, designed the evaluation framework, implemented the analysis pipeline, and wrote the manuscript.

## Data availability

Processed result tables and figures are available at https://github.com/AsiaeeLab/tissue-aware-drug-response. The underlying CCLE/DepMap molecular data and GDSC drug response data are available from their original sources.

