## Supplementary Material for "Tissue-aware elastic net decomposition reveals shared and lineage-specific drug response biomarkers"

This document contains supplementary tables and figures referenced in the main manuscript. Throughout, items are numbered S1, S2, ...; references to main-text Figures 1–4 and Tables 1–4 refer to the items in the main manuscript.

#### Supplementary Tables

Table S1: **Aggregate lineage breadth of stably-selected gene blocks.** For each method's stable feature set ($\geq$ 50% bootstrap selection), each selected gene–drug pair is mapped to its tissue-level expression-specificity score $\tau$ (defined in Methods), where lower values indicate broader expression across the 31 GDSC lineages and higher values indicate lineage restriction. "Broadly expressed" is the fraction of gene blocks in the lowest $\tau$ quartile of the EN reference distribution; "lineage-restricted" is the fraction in the highest quartile. The DSEN shared block is modestly broader than the DSEN tissue block (median $\tau$ 0.136 vs. 0.147; Wilcoxon $p = 0.040$), but the shift is small in absolute terms, supporting a careful rather than absolute pan-cancer-vs-lineage interpretation.

| Block | Gene blocks | Unique genes | Median $\tau$ | Broadly expressed | Lineage-restricted |
|---|---|---|---|---|---|
| EN | 6,037 | 2,377 | 0.139 | 16.0% | 41.3% |
| DSEN shared | 2,296 | 1,211 | 0.136 | 15.9% | 42.5% |
| DSEN tissue | 7,408 | 2,647 | 0.147 | 10.7% | 42.3% |

Table S2: **Leave-skin-out generalization for the 11 BRAF/MEK-pathway compounds.** For each strict BRAF/MEK inhibitor (including replicates), we held out all skin cell lines, retrained DSEN on the remaining tissues, and used only the shared coefficient block to predict the held-out skin samples. Improvement is the relative reduction in MSE over the elastic-net baseline trained under the same protocol (positive = lower MSE). DSEN's shared block loses on 10 of 11 cases (median −19.2%, mean −21.8%), identifying the skin tissue block as the lineage-specific component that fails to transfer for this drug class.

| Drug | Held-out cells | MSE (EN) | MSE (DSEN shared) | Improvement (%) |
|---|---|---|---|---|
| Dabrafenib | 47 | 0.2141 | 0.3549 | −65.7 |
| Selumetinib (replicate) | 53 | 0.1190 | 0.1744 | −46.5 |
| Refametinib (replicate) | 44 | 0.1069 | 0.1412 | −32.0 |
| Refametinib | 50 | 0.1201 | 0.1503 | −25.1 |
| PLX-4720 | 46 | 0.0770 | 0.0921 | −19.6 |
| CI-1040 | 50 | 0.0853 | 0.1017 | −19.2 |
| PD0325901 | 44 | 0.1240 | 0.1421 | −14.5 |
| PLX-4720 (replicate) | 56 | 0.0619 | 0.0697 | −12.7 |
| Trametinib | 51 | 0.2457 | 0.2542 | −3.4 |
| Selumetinib | 43 | 0.0729 | 0.0747 | −2.5 |
| SB590885 | 42 | 0.0726 | 0.0711 | +2.1 |

Table S3: **Leave-tissue-out generalization, per tissue.** For each tissue, we list the number of drugs for which the tissue was held out, the total number of (drug, held-out tissue) predictions made by DSEN's shared coefficient block, the tissue-level win rate (fraction of predictions with DSEN shared MSE below the EN baseline), and the mean and median relative MSE improvement (positive = DSEN better). Tissues are ordered by mean improvement. Aggregated across 224 drugs with $\geq 5$ tissues, DSEN's shared block won on 58.8% of 2,800 tissue-holdout predictions with a mean per-drug improvement of 1.8%; 149 of 224 drugs (66.5%) had DSEN winning on more than half their tissues.

| Held-out tissue | Drugs | Predictions | Win rate (%) | Mean impr. (%) | Median impr. (%) |
|---|---|---|---|---|---|
| Large intestine | 211 | 224 | 77.2 | +17.06 | +10.18 |
| Upper aerodigestive tract | 204 | 217 | 65.4 | +6.48 | +4.20 |
| Skin | 211 | 224 | 65.6 | +4.87 | +3.41 |
| Central nervous system | 211 | 224 | 63.4 | +4.72 | +2.85 |
| Kidney | 119 | 121 | 58.7 | +3.11 | +1.97 |
| Autonomic ganglia | 198 | 208 | 58.7 | +2.26 | +1.46 |
| Lung | 211 | 224 | 67.0 | +2.18 | +1.88 |
| Ovary | 207 | 220 | 55.5 | +1.68 | +0.83 |
| Haematopoietic/lymphoid | 211 | 224 | 54.9 | +1.24 | +0.42 |
| Stomach | 46 | 47 | 57.4 | +0.92 | +0.71 |
| Breast | 210 | 223 | 48.4 | +0.39 | −0.05 |
| Pancreas | 193 | 203 | 56.2 | +0.37 | +0.21 |
| Oesophagus | 204 | 217 | 48.4 | −1.41 | −0.78 |
| Bone | 211 | 224 | 44.6 | −2.15 | −0.99 |

### Supplementary Figures

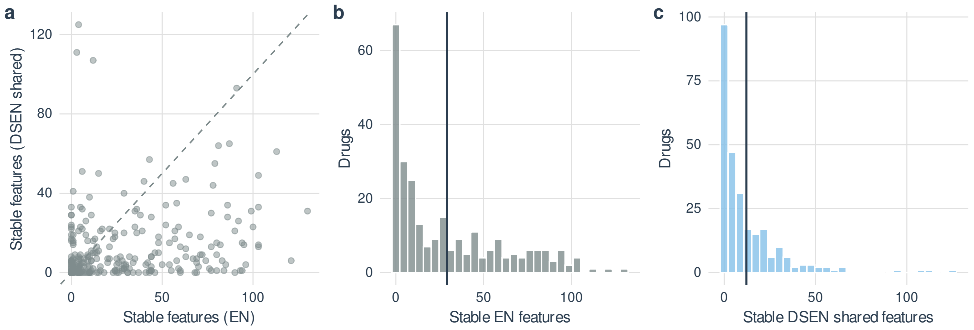


Figure S1: **DSEN yields sparser shared feature sets than standard elastic net.** **(a)** Scatter of stable feature counts (selected in ≥50% of 200 bootstrap resamples) for elastic net versus the DSEN shared block; most drugs fall below the diagonal. **(b)** Distribution of EN stable feature counts (mean 29). **(c)** Distribution of DSEN shared stable feature counts (mean 12.2). Overall, 63.8% of drugs have sparser DSEN shared blocks than the standard elastic-net baseline.
